# A Canonical Microcircuit for Estimating Excitation/Inhibition (E/I) Balance

**DOI:** 10.1101/2025.07.10.664116

**Authors:** Daniel J. Hauke, Julia Rodriguez-Sanchez, Hope Oloye, Lioba C. S. Berndt, Dimitris Pinotsis, Karl J. Friston, Daniel H. Mathalon, Rick A. Adams

**Affiliations:** Hawkes Institute, University College London, UK; School of Health and Medical Sciences, City St George’s, University of London, UK; Department of Psychology, Faculty of Health and Life Sciences, University of Exeter, UK; Department of Psychology and Neuroscience, City St George’s, University of London, UK; Department of Imaging Neuroscience, Queen Square Institute of Neurology, University College London, UK; Department of Psychiatry & Behavioral Sciences, University of California, San Francisco, CA, USA; Mental Health Service, San Francisco Veterans Affairs Health Care System, San Francisco, CA, USA; Institute of Cognitive Neuroscience, University College London, UK

**Keywords:** E/I balance, dynamic causal modelling, predictive coding, biophysical modelling, EEG, MEG

## Abstract

Excitation/inhibition (E/I) balance is crucial for maintaining healthy brain function and can be disrupted in various neurological and psychiatric disorders. Despite its importance, there are few tools to study E/I balance non-invasively in humans. Here, we propose a canonical microcircuit model to estimate E/I balance from non-invasive magneto- and electroencephalography (M/EEG) recordings by parameterising global pyramidal and inhibitory cell excitability. We first establish that E/I parameters are identifiable and recoverable. We then explore the effects of these new parameters and their interaction with other parameters in a series of simulations. To highlight the clinical relevance of this new model, we simulate changes in E/I balance and their impact on event-related potentials (ERPs) derived from paired-click, passive and active oddball paradigms, which are among the most robust clinical biomarkers of schizophrenia. Our simulations show that a loss of pyramidal cell excitability can explain reduced ERP amplitudes across all three paradigms, mirroring empirical findings in schizophrenia. This method may serve as a computational assay for estimating synaptopathy and E/I balance from non-invasive M/EEG recordings across various clinical conditions thereby advancing efforts to develop personalised interventions to restore E/I balance.

## Introduction

Our brain relies on a fine-tuned balance between excitation (E) and inhibition (I), referred to as E/I balance, to maintain healthy function. Even small increases in E/I can lead to runaway excitation, as observed in epileptic seizures (Chagnac-Amitai, & Connors, 1989). Conversely, excessively increased inhibition can lead to sedation - for example, after alcohol intake (Ticku, 1990) - or even to loss of consciousness (Eisen et al., 2024). E/I balance can be manipulated pharmacologically in various ways through drugs acting on glutamatergic and gamma-aminobutyric-acid (GABA)-ergic targets, but also through psychotomimetics like ketamine (Fagerholm et al., 2021; Umbricht et al., 2002; Javitt et al., 1996) and possibly psychedelics like psilocybin (Leptourgos et al., 2022) and lysergic acid diethylamide (Bedford et al., 2023; Hauke et al., 2025; Leptourgos et al., 2022).

Disturbances of E/I balance have featured in aetiological theories of numerous neurological and psychiatric disorders like epilepsy (Fritschy, 2001), autism (Rubenstein, & Merzenich, 2003), depression (Hu et al., 2023), and schizophrenia (Krystal et al., 2017; Sohal, & Rubenstein, 2019; Yizhar et al., 2011). In the case of schizophrenia, for instance, multiple research lines support a role for E/I imbalance, including genetic evidence implicating α-amino-3-hydroxy-5-methyl-4- isoxazolepropionic acid (AMPA) and N-methyl-D-aspartate (NMDA) receptors (Singh et al., 2022; Trubetskoy et al., 2022), and postmortem studies that suggest reduced dendritic spine density and interneuron deficits (Glantz, & Lewis, 2000; Lewis, Hashimoto, & Volk, 2005). Additionally, electrophysiological signals measured with electroencephalography (EEG) and magnetoencephalography (MEG) are changed in individuals at clinical high risk for psychosis and patients with schizophrenia. This includes reduced gamma-band activity (Grent-’t-Jong et al., 2023), mismatch negativity (Erickson et al., 2016; Garrido et al., 2009a; Hamilton et al., 2020; Hamilton & Mathalon, 2024; Umbricht & Krijes, 2005) and P300 in oddball paradigms (Bramon et al., 2004; Jeon et al., 2003; Hamilton et al., 2020; Hamilton & Mathalon, 2024), and reduced event-related potentials (ERPs) in paired-click paradigms (Turetsky et al., 2008; Shen et al., 2020; Rosburg, 2018). There are pharmacological studies that reproduce ERP reductions observed in schizophrenia with ketamine and GABAergic drugs suggesting potential links with E/I balance (Gunduz-Bruce et al., 2012; Javitt et al., 1996; Hamilton et al., 2018; Mathalon et al., 2014; Oranje et al., 2000; Plourde et al., 1997; Rosburg & Kreitschmann-Andermahr, 2016; Schwertner et al. 2018; Umbricht et al., 2002; Watson et al., 2009). However, the field needs new methods to estimate E/I balance *in vivo* from non-invasive recordings to identify the precise nature of circuit dysfunction underlying these M/EEG biomarkers.

Here, we propose a canonical microcircuit model specifically designed to estimate E and I cell function from non-invasive M/EEG recordings with dynamic causal modelling. We simulate how changes in E/I parameters can explain ERP alterations observed in schizophrenia to highlight their predictive validity, and we systematically assess how these parameters interact with each other through a series of simulations. The aims of this paper are threefold: 1) to introduce this new method for studying E/I balance, 2) to demonstrate the model’s utility for studying neuropsychiatric disorders using schizophrenia as an example, and 3) to provide a comprehensive resource for the scientific community by making the modelling code publicly available along with empirically-informed priors, for future investigations of E/I balance across various clinical conditions.

## Methods

### Canonical Microcircuit for estimating E/I balance

Our model is grounded in the dynamic causal modelling (DCM) framework. This state-space modelling framework was originally introduced to model fMRI data (Friston, 2002; Friston, Harrison, & Penny, 2003) and later extended to M/EEG (David & Friston, 2003; David et al., 2006; Kiebel, David, & Friston, 2006; Kiebel et al., 2008) based on seminal work from Jansen and Rit (Jansen & Rit, 1995). This modelling approach can be concisely summarised by two sets of equations:

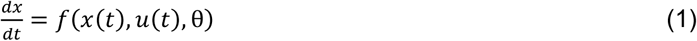

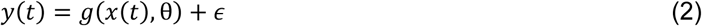

The state equation (Eq. 1) describes how hidden states *x* (in this case the voltage and current of neuronal populations) evolve in time as a function of the states *x* at time *t*, a thalamic input into the system *u* at time *t* and model parameters θ (e.g., expressing synaptic connectivity between neuronal populations). The observer equation (Eq. 2) describes how changes in the hidden states *x* are mapped onto the measured signal *y* at time *t* (e.g., an EEG recording) through an observation model with parameters θ and a normally distributed observation error *ϵ*.

DCM for EEG comes in two forms: conductance-based and convolution-based models. The former models the membrane conductance of each neuronal population and specific ion channels (GABA, AMPA and NMDA receptors) based on Hodgkin and Huxley’s model of the squid giant axon (Hodgkin & Huxley, 1952) scaled from single cells to neuronal populations with the Morris & Lecar formalism (Morris & Lecar, 1981; see Pinotsis et al., 2013, and Pereira et al., 2021, for an overview). Conversely, convolution-based DCM models are based on Jansen & Rit (1995) and model the conversion of incoming spikes to postsynaptic potentials (PSPs) through convolution with a synaptic response kernel (David & Friston, 2003; David et al., 2006; Kiebel, David, & Friston, 2006; Kiebel et al., 2008; Moran et al., 2013; Pinotsis et al., 2016). The E/I model we propose generalises across both convolution-based and conductance-based models, but we focus on the convolution-based model to introduce the E/I parameters. Next, we will unpack the modelling approach across different spatial scales.

#### Microscale Model: Neuronal Population

At the microscale, we model a neuronal population through two basic operations (Figure 1). First, at dendritic spines, incoming spikes are converted into PSPs through convolution with a synaptic response function:

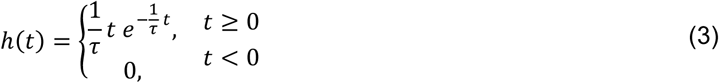

where τ is the synaptic time constant, with larger *τ* leading to more prolonged responses in time (Figure 1).

**Figure 1.**
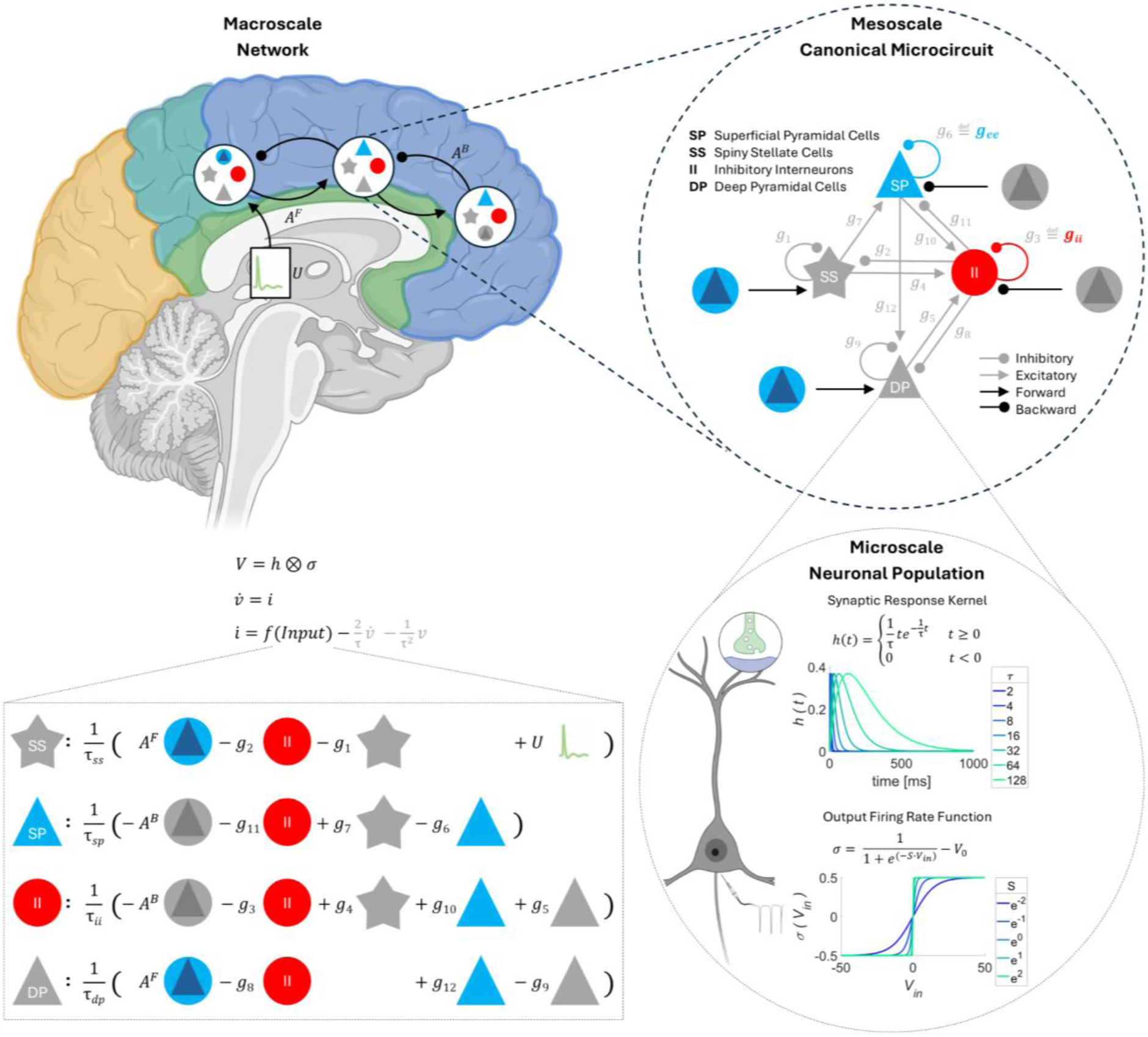
Model Overview. At the macroscale, the model requires defining a network of relevant sources that receive a thalamic input *U* and are connected via extrinsic forward *A*^*F*^ and backward *A*^*B*^. At the mesoscale, this model extends the canonical microcircuit for predictive coding (Bastos et al., 2012; Pinotsis et al., 2013), modelling each source as a microcircuit with four neuronal populations: E cells or superficial pyramidal cells (**SP**), spiny stellate cells (**SS**), I cells or inhibitory interneurons (**II**) and deep pyramidal cells (**DP**). These populations are connected via intrinsic connections *g* according to Shaw et al. (2017). To estimate E/I balance, we parameterise the global (across-region) excitability of excitatory pyramidal *g*_*ee*_ and inhibitory cells *g*_*ii*_ modelled as (inhibitory) self-connections. At the microscale each neuronal population is modelled through two basic operations 1) a convolution with a synaptic response kernel and 2) a sigmoid function to map postsynaptic potentials onto an output firing rate. The figure was created in BioRender (Hauke, 2025).

Secondly, PSPs *V*_*in*_ are converted into an average spike rate σ at the axon hillock using a sigmoid function:

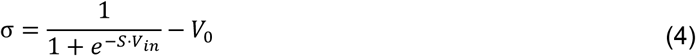

where *V*_0_ indicates the baseline firing rate, *V*_*in*_ is average membrane potential of the neuronal population, and *S* parametrizes the slope of the sigmoid function and captures the dispersion of firing thresholds within this neuronal population, with smaller values of *S* resulting in a shallower slope (i.e., more dispersed thresholds in the population, Figure 1). Each population can be modelled through the convolution of these two functions:

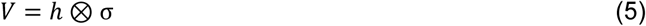

which results in a system of two coupled ordinary differential equations for each neuronal population, the first describing the change in current and the second describing the change in voltage:

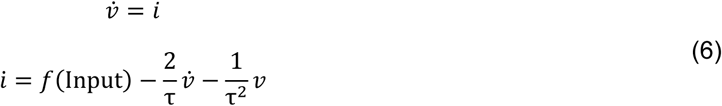

The input is determined by intrinsic (within-source) and extrinsic (between-source) connectivity.

#### Mesoscale Model: Canonical Microcircuit

At the mesoscale, we model a cortical column using the canonical microcircuit for predictive coding comprising inhibitory interneurons, spiny stellate cells, as well as superficial and deep pyramidal cells, based on previous work (Douglas & Martin, 1991; Haeusler & Maass, 2006; Bastos et al., 2012; Pinotsis et al., 2013). We connect these neuronal populations in the same cortical source based on Shaw et al. (2017; Figure 1).

Importantly, we parameterise the self-connection of E (superficial pyramidal cells) and I cells (inhibitory interneurons), which we refer to as *g*_*ee*_ and *g*_*ii*_, where the first subscript indicates the origin of the connection (*e* for the excitatory and *i* for the inhibitory population) and the second subscript specifies the target population. These parameters determine the excitability of E and I cells and will change their average output firing rate. In this study, we operationalise E/I ratio as the ratio of average output firing rate of E and I cells that can be manipulated through changing these two parameters.

Increasing *g*_*ee*_, for example, corresponds to increasing the connection strength of the (inhibitory) self-connection, which models a loss of E cell excitability (or reduced E/I ratio). Conversely, increasing *g*_*ii*_ models a loss of I cell excitability resulting in net disinhibition of the microcircuit. Both self-connections are always inhibitory to ensure that the model generates stable dynamics. We can generalise this approach to the conductance-based model by parameterising the (self-inhibitory) GABAergic connections of the same neuronal populations.

We make two important modelling choices regarding these E/I parameters: 1) we constrain these parameters to be the same across all modelled regions in order to ensure an efficient global estimate of E and I cell excitability and 2) we allow them to be both general and condition-specific, expressing the average E/I balance across all conditions and the difference in E/I balance between conditions, respectively.

First, whilst there may be regional variation in E and I, we assume that genetic predisposition to altered E/I is generalised across cortex than confined to specific areas (e.g., based on Fish et al., 2021; Osimo et al., 2019; Shenton et al., 2001 in the case of schizophrenia-related changes). This is reflected in our cortex-wide parameter with the added potential benefit of using all the EEG data to estimate it. Note, however, that this assumption can be relaxed if one wishes to model gradients of E/I balance across the brain.

Second, we introduce these E and I parameters both as condition-specific parameters (capturing condition-specific rebalancing of E/I ratio in response to experimental inputs), referred to as *B*^*g_ee_*^ and *B*^*g_ii_*^ in the remainder of the paper, and condition-independent parameters expressing the average E/I balance, referred to as *g*_*ee*_ and *g*_*ii*_. This is motivated by the notion that E/I changes are adaptive and may well have more impact on some EEG conditions, e.g. those that typically generate a much larger response (like oddballs).

#### Macroscale Model: Network

Lastly, we connect different sources with excitatory forward connections *A*^*F*^ that originate from superficial pyramidal cells in one source and project to spiny stellate cells and deep pyramidal cells of another source, and inhibitory backward connections *A*^*B*^ which originate from deep pyramidal cell populations and project to superficial pyramidal cells and inhibitory interneurons (Bastos et al., 2012; Pinotsis et al., 2013). This results in the following system of coupled differential equations for each source. Note that these coupled equations have the form of Eq. (6), but for consistency with previous work and the provided code, we denote all hidden states (voltage and current) with *x*. For spiny stellate cells, we have:

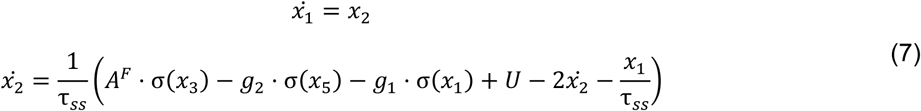

For E cells (superficial pyramidal cells), we have:

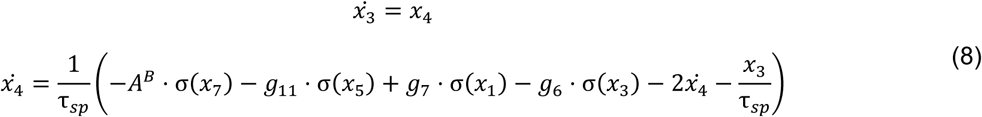

For I cells (inhibitory interneurons), we have:

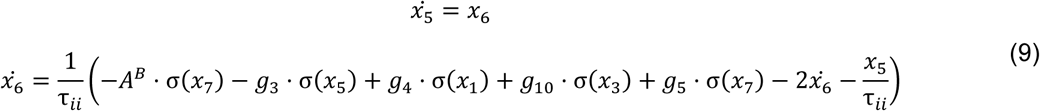

And lastly for deep pyramidal cells, we have:

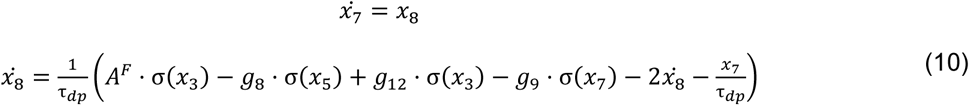

We implemented this model using SPM12 (version 7771, Wellcome Trust Centre for Neuroimaging, London, UK, https://github.com/spm/spm12/releases/tag/r7771) in MATLAB (version R2020b; The MathWorks Inc., Natick, MA, USA; https://www.mathworks.com). The code for both the convolution- and conductance-based E/I models can be found on GitHub (https://github.com/daniel-hauke/dcm_ei).

### Simulations

To illustrate the effects of the E and I parameters, we conduct a series of simulations based on parameter estimates from data collected from a large number of healthy volunteers during three different paradigms as part of the Bipolar and Schizophrenia Network for Intermediate Phenotypes (B-SNIP; Tamminga et al., 2013; Clementz et al., 2016) and the second North American Prodrome Longitudinal Study (NAPLS2; Addington et al., 2012; Hamilton et al., 2019; Hamilton et al., 2022). We briefly recapitulate the most pertinent information about the paradigms and preprocessing below. For more detailed information on task and preprocessing, please see the referenced studies.

#### Participants

We analysed EEG data from *n*=266 (paired-click paradigm), *n*=241 (passive oddball paradigm), and *n*=237 (active oddball paradigm) healthy participants. The EEG setup and preprocessing is reported in the Supplement. To provide priors for future analyses – that can be used beyond investigations focused on schizophrenia – we computed the grand mean data from healthy control participants. We ground our simulations in these denoised grand mean ‘participants’ to illustrate the effects of changing E and I parameters.

#### Tasks

##### Paired-click paradigm

Participants were presented with 150 binaural broadband auditory click pairs (4ms, 75dB, inter-click interval: 500ms, average interstimulus interval: 9.5s; Hamm et al., 2014).

##### Passive oddball paradigm

Participants performed a visual oddball primary task while being presented with 1794 tones that were either standard tones (85%,633Hz, 50ms), duration deviants (5%, 633Hz, 100ms), pitch deviants (5%, 1000Hz, 50ms) or duration+pitch double-deviants (5%, 1000Hz, 100ms). Tone length included 5ms rise/fall time. Subjects were told to ignore the tones and focus on the visual task. Here, we focused on modelling the double-deviant tone as it showed the largest clinical effects in terms of predicting conversion to psychosis in individuals at clinical high risk for psychosis (Hamilton et al., 2022).

##### Active oddball paradigm

In the active oddball paradigm, participants were presented with 450 stimuli including 80% standard tones (500Hz, 50ms with 5ms rise/fall time),10% target tones (1000Hz, 50 ms, 5ms rise/fall time) and 10% novel sounds with a stimulus onset asynchrony of 1250ms (Hamilton et al., 2019). Novel sounds were various natural and man-made sounds (Friedman, Simpson, & Hamberger, 1993). Participants were asked to respond to target tones by pressing a button.

#### Networks

We defined three task-specific networks to be modelled with our E/I model.

##### Paired-click paradigm

Since we are unaware of previous studies that used DCM to model paired-click responses, we defined a task network based on previous literature (Bak et al., 2011; Nakagawa et al., 2014) comprising left (MNI: [-53 -20 13]) and right superior temporal gyrus (STG; [49 -10 10]) as auditory input regions, left ([-36 23 3]) and right insular cortex (Ins; [36 23 3]), and left ([-1 -20 67]) and right medial frontal gyrus (MFG; [1 -20 67]). To confirm that these sources explain sufficient variance in the data, we conducted a multiple sparse prior (MSP) source analysis with 16mm spheres drawn around these coordinates and compared it to an (unconstrained) independent and identically distributed (IID) source analysis in SPM. These six sources explained 95.06% of the variance versus 97.18% in the IID case. Formal model comparison based on the model evidence suggested that there was strong evidence that the 6-region source reconstruction was the better model (log Bayes factor: 23.6; Kass & Raftery, 1995).

##### Passive oddball paradigm

We modelled the passive oddball paradigm with a well-established 6-region network comprising bilateral primary auditory cortex (A1; left: [-42 -22 7], right: [46 14 8]) as input regions, STG (left: [-61 - 32 8], right: [59 -25 8]) and inferior frontal gyrus (IFG; left: [-46 20 8], right: [46 20 8]) based on extensive previous work (Doeller et al., 2003; Garrido et al., 2007; Garrido et al., 2009a; Molholm et al., 2005; Opitz et al., 2002; Rademacher et al., 2001; Rinne et al., 2000) including various DCM studies (Adams et al., 2022; Garrido et al., 2009b; Schmidt et al., 2013).

##### Active oddball paradigm

We defined a 6-region network for the active oddball paradigm based on a previous meta-analysis of 75 oddball paradigm PET/fMRI studies (Kim, 2014). We chose STG as input regions (left: [-61 -32 8], right: [59 -25 8]) using the same coordinates as for the passive oddball task network and confirmed that they fell into the brain region activated during the active oddball task (Kim, 2014). Importantly, in contrast to the passive oddball paradigm, the active oddball paradigm requires participants to maintain top-down attentional control towards the deviant tones to respond with button presses (Debener et al., 2002). We thus modelled bilateral inferior frontal junction (IFJ; left: [-56 7 29], right: [50 8 30]) and intraparietal sulcus (IPS; left: [-33 -42 64], right: [33 -42 64]) regions of the dorsal attention network based on Kim (2014). We found that this 6-region model explained 94.31% of the variance versus 95.14% of an IID source reconstruction and was strongly favoured in model comparison (log Bayes factor: 49.2).

#### Integration

We integrated the convolution-based model with a Euler integrator (Schöbi et al., 2021), which we found to have the best accuracy-runtime trade-off for our purposes. We also provide code that works with the default SPM12 integrators and an implementation using MATLABs dde23 integrator for benchmarking the conductance-based model (Supplement).

#### Priors

We used the default SPM12 priors with the following changes. In the source model, we implemented equality constraints on transmission delays (extrinsic and intrinsic) and intrinsic connectivity parameters across regions to drastically reduce the number of free parameters by 40 (20→2 delay and 24→2 intrinsic connectivity parameters) for a six-region model. We used the equivalent current dipole (ECD) forward model and assumed that only E cells (superficial pyramidal cells) contributed to the EEG signal (fixing an additional 2 parameters), although note that this assumption can be easily relaxed.

The neuronal time constants *τ* of the original canonical microcircuit model were selected to model gamma oscillations (Pinotsis et al., 2013). We found that, under the new integration schemes, the default priors on *τ* were not suitable for generating the oscillations that we wished to model – especially for the active oddball task, where the characteristic responses are on slower (theta) time scales. In simulations, we also observed that the slope of the sigmoid output firing rate function *S* — parametrising the dispersion of firing thresholds or depolarisation of a neuronal population (Marreiros et al., 2008) — was crucial in ensuring that the model generated dampened as opposed to ongoing or increasing oscillations. We thus performed a grid search over different combinations of the four time constants and *S* spanning the following grids *τ*_*ss*_: [2 16 32 64] ms, *τ*_*ii*_ : [2 16 32] ms, *τ*_*sp*_: [2 16 32 64 128] ms, *τ_dp_*: [2 16 32 64 128] ms and *S*: [-2 -1 0 1 2]. We identified the best model based on the free energy approximation to the log model evidence, while constraining the solutions to physiological plausible parameter ranges, which we defined as [1 60] ms for spiny stellate cells (Markram et al., 2015; Ramaswamy et al., 2015; Stern, Edwards, & Sagman, 1992; Sun, Huguenard, & Prince, 2006), [1 30] ms for inhibitory interneurons (Bartos et al., 2001; Hájos, & Mody, 1997) and [1 200] ms for pyramidal cells. The *τ* range for pyramidal cells was motivated by evidence that NMDA receptors on these neurons can have time constants of around 40-200 ms (Wang et al., 2008; Zhang et al., 2016), and these receptors may be captured implicitly in the convolution-based model. All priors are summarised in Table 1.

**Table 1.** Parameter Priors. Note, that the grid search identified different priors for the slope of the sigmoid function *S* and the synaptic time constants *T* for different paradigms (cf. *Priors* section). The optimal values are determined by the overall duration of the time window that is modelled and the dominant frequencies; for example, more alpha in paired-click and passive oddball paradigms compared to more theta in the active oddball paradigm. The input parameters were selected to ensure that the input peak coincides with the first dominant peak of the event-related potential of each paradigm. For the motivation of the three different networks (**Lpos**), please see the *Networks* section. **DP** deep pyramidal cells. **II** inhibitory interneurons. **SP** superficial pyramidal cells. **SS** spiny stellate cells.

| Parameter | Description | Prior Mean | Prior Precision |
| --- | --- | --- | --- |
| **Neuronal Source Model $f(x)$** |  |  |  |
| **A{1}** | Average extrinsic connectivity (forward, SP to SS) | 0 | 1/16 |
| **A{2}** | Average extrinsic connectivity (forward, SP to DP) | 0 | 1/16 |
| **A{3}** | Average extrinsic connectivity (backward, DP to SP) | 0 | 1/16 |
| **A{4}** | Average extrinsic connectivity (backward, DP to II) | 0 | 1/16 |
| **B** | Condition-specific extrinsic connectivity | 0 | 1/8 |
| **C** | Input Strength | 0 | 1/32 |
| **G** | Average intrinsic connectivity [$g_{ee}$ $g_{ii}$] | 0 | 1/32 |
| **B<sup>G</sup>** | Condition-specific intrinsic connectivity [$B^{gee}$ $B^{gii}$] | 0 | 1/8 |
| **T** | Synaptic time constant [SS SP II DP] | Paired Click: [16 16 2 16] ms<br>Passive oddball: [16 32 2 2] ms<br>Active oddball: [16 128 2 2] ms | 1/32 |
| **D** | Neuronal delays (all sources) [intrinsic extrinsic] | [1 8] ms | 1/64 |
| **S** | Slope of output firing rate function | Paired Click: -2<br>Passive oddball: -1<br>Active oddball: -1 | 1/64 |
| **R** | Stimulus [onset dispersion] | Paired Click: [50 30] ms<br>Passive oddball: [68 16] ms<br>Active oddball: [100 16] ms | 1/1024 |
| **Forward Model $g(x)$** |  |  |  |
| **Lpos** | Source location [MNI coordinates] | Paired Click: [-53 -20 13;<br>49 -10 10;<br>-36 23 3;<br>36 23 3;<br>-1 -20 67;<br>1 -20 67]<br>Passive oddball: [-42 -22 7;<br>46 -14 8;<br>-61 -32 8;<br>59 -25 8;<br>-46 20 8;<br>46 20 8]<br>Active oddball: [-61 -32 8;<br>59 -25 8;<br>-56 7 29;<br>50 8 30;<br>-33 -42 64;<br>33 -42 64] | 0 |
| **L** | Dipole orientation | 0 | 64 |
| **J** | Neuronal populations contributing to the EEG:<br>[SS SP II DP] | [0 1 0 0] | 0 |

#### Model diagnostics

To assess model fit (i.e., accuracy), we computed the Pearson correlation between the responses predicted by the model and the measured EEG responses. To assess parameter identifiability, we fit the model to 100 randomly selected healthy participants using the posteriors from the grand mean as empirical priors. We report the Pearson correlation between E/I parameters across participants to test whether parameters are identifiable (i.e., their posterior dependencies or correlations are small). In line with previous work (Hauke et al., 2024), we consider *r*<|0.6| to be an acceptable correlation, suggesting that the parameters mediate distinct processes. To measure parameter recovery, we (i) simulated synthetic data using the posterior parameters from models fit to 100 randomly selected participants, (ii) added empirical levels of noise, estimated from the data (i.e., using the posterior estimate saved in DCM.Ce), and (iii) reinverted the models on the simulated data. We report intraclass correlation coefficients (ICC), as recommended previously (Karvelis, Paulus & Diaconescu, 2023; Karvelis et al., 2024), and consider recovery to be poor (<0.4), fair (0.4–0.59), good (0.6–0.74), or excellent (>0.75) as suggested by Fleiss (2011). We also report Pearson correlations between simulated and recovered parameters in line with previous work (Hauke et al., 2022; 2024).

#### Parameter sensitivity analysis

To illustrate the effects of the E and I parameters, we simulated ERPs starting with their posterior and incrementally increased or decreased the parameters over the interval [-0.5 0.5] in steps of 0.125, which corresponds to scaling the connectivity strength of the self-connection by ±50%, while keeping all other parameters fixed to their posterior estimate. We also varied multiple parameters together. To simplify the visualisation for multidimensional simulations, we display the amplitude of characteristic ERP components. Specifically, we defined the P2 component as the most positive deflection in Cz between 120-250ms in the paired-click paradigm, the MMN, as the most negative deflection in Fz between 150-250ms in the passive oddball paradigm, and the P3b component as the most positive deflection in Pz between 250-600ms in the active oddball paradigm.

## Results

### Model diagnostics

The model fit all paradigms remarkably well. Specifically, it predicted responses in the paired stimulus paradigm with an average *r*=0.95 (tone 1: *r*=0.95; tone 2: *r*=0.96; Figure 2a), in the passive oddball paradigm with *r*=0.89 (standard: *r*=0.85; deviant: *r*=0.94; Figure 2d) and the active oddball paradigm with *r*=0.84 (standard: *r*=0.71; target: *r*=0.97; Figure 2g). Note that ERPs tend to be much larger in the deviant and target conditions of the oddball paradigm compared to the standard tone condition, while the noise is comparable between conditions. This results in reduced SNR in the standard condition and likely explains why the model shows reduced fits for this condition. We observed good parameter identifiability (all *r*<|0.6| across all paradigms; Figure 2b, 2e and 2h). Parameter recovery of *B*^*g_ee_*^ was excellent, while *B*^*g_ii_*^ recovery was fair across all three paradigms (Figure 2c, 2f, 2i).

**Figure 2.**
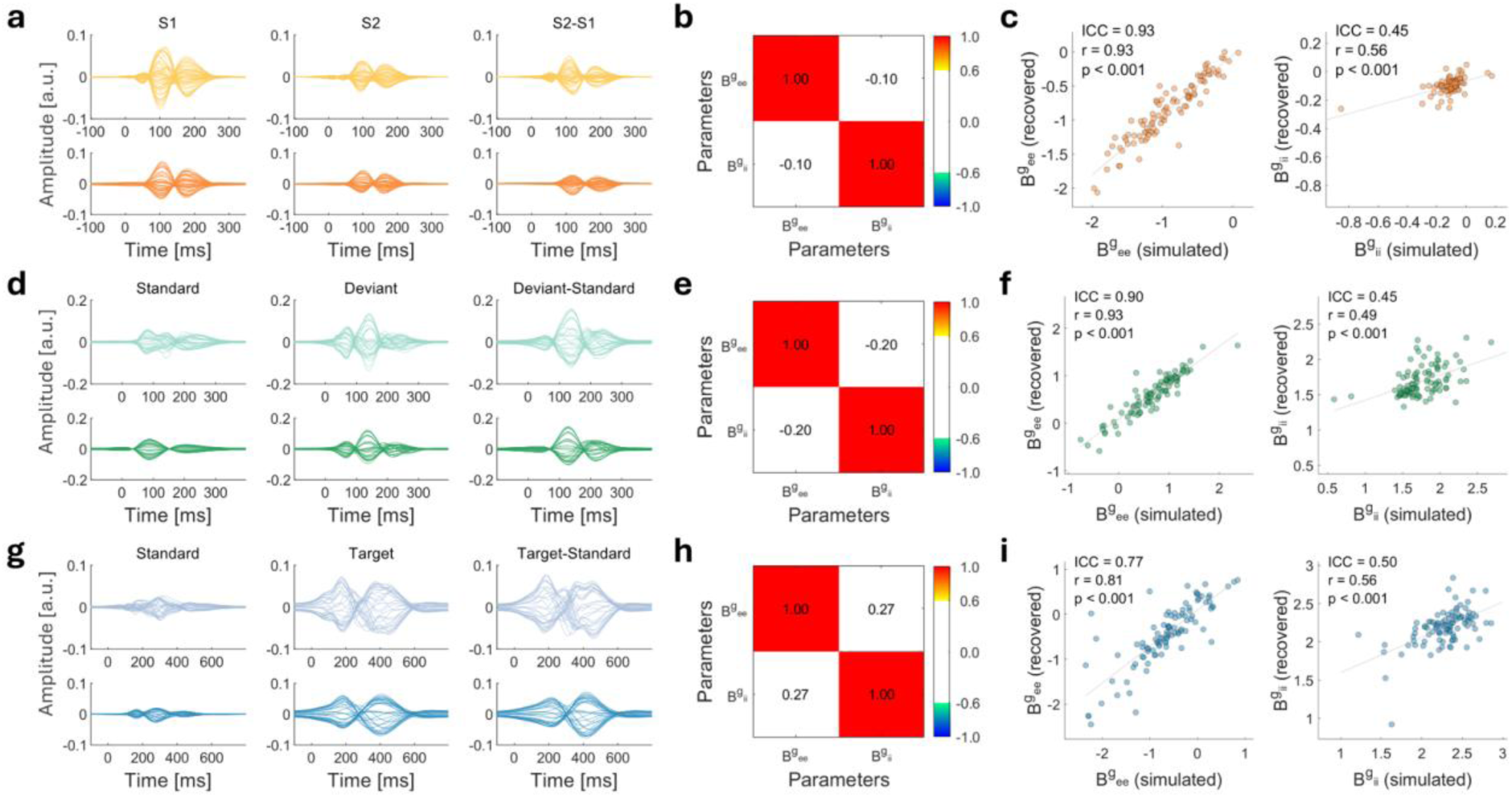
Model diagnostics. Shown are observed responses (lighter colours) and responses predicted by the model (darker colours) for the paired-click (**a**), passive oddball (**d**) or active oddball paradigm (**g**). Correlation matrices with between-subject parameter correlations for 100 randomly selected participants are displayed to illustrate parameter identifiability for the paired-click (**b**), passive oddball (**e**) or active oddball paradigm (**h**). We interpret parameter correlations *r*<|0.6| as indicative of acceptable parameter identifiability (i.e., limited conditional dependency) in line with Hauke et al. (2024). Parameter recovery is shown for the paired-click (**c**), passive oddball (**f**) or active oddball paradigm (**i**). To measure parameter recovery, we simulated synthetic data using the posterior parameters (x-axis) from models fit to 100 randomly selected participants adding empirical levels of noise, which were estimated from the data, and reinverted the models on the simulated data to obtain recovered parameters (y-axis). We consider intraclass correlation coefficients (**ICC**) to be indicative of poor (<0.4), fair (0.4–0.59), good (0.6–0.74), or excellent (>0.75) recovery in line with Fleiss (2011).

### Posterior parameter estimates

Posterior parameter estimates are reported in Supplementary Tables 1-3 and provided in SPM12 DCM format on GitHub (https://github.com/daniel-hauke/dcm_ei). We recommend using these parameters as empirical priors for future DCM analyses with this model and these paradigms.

### Simulating changes in E and I cell excitability

To illustrate the validity of the model, we simulated a loss of condition-specific E or I cell excitability across all three paradigms and compared those results to known effects in the schizophrenia literature (Figure 3). We found that increasing *B*^*g_ee_*^ to simulate a loss of pyramidal cell excitability reduced P1 and N2 components of the difference waveform in the paired click paradigm by reducing the response to the first tone and increasing the response to the second tone (Figure 3a), which may explain reduced S2/S1 ratios observed in patients (Turetsky et al., 2008; Shen et al., 2020; Rosburg, 2018). Another finding in the schizophrenia literature is that patients show reduced responses to both tones (Turetsky et al., 2008), which can be simulated with our model by increasing *g*_*ee*_ (reducing average excitability across both conditions, Figure S1). Interestingly, simulating a loss of inhibitory cell excitability by increasing *B*^*g_ii_*^ had no effect on the waveform, suggesting that this parameter alone cannot explain the changes observed in schizophrenia (Figure 1b).

**Figure 3.**
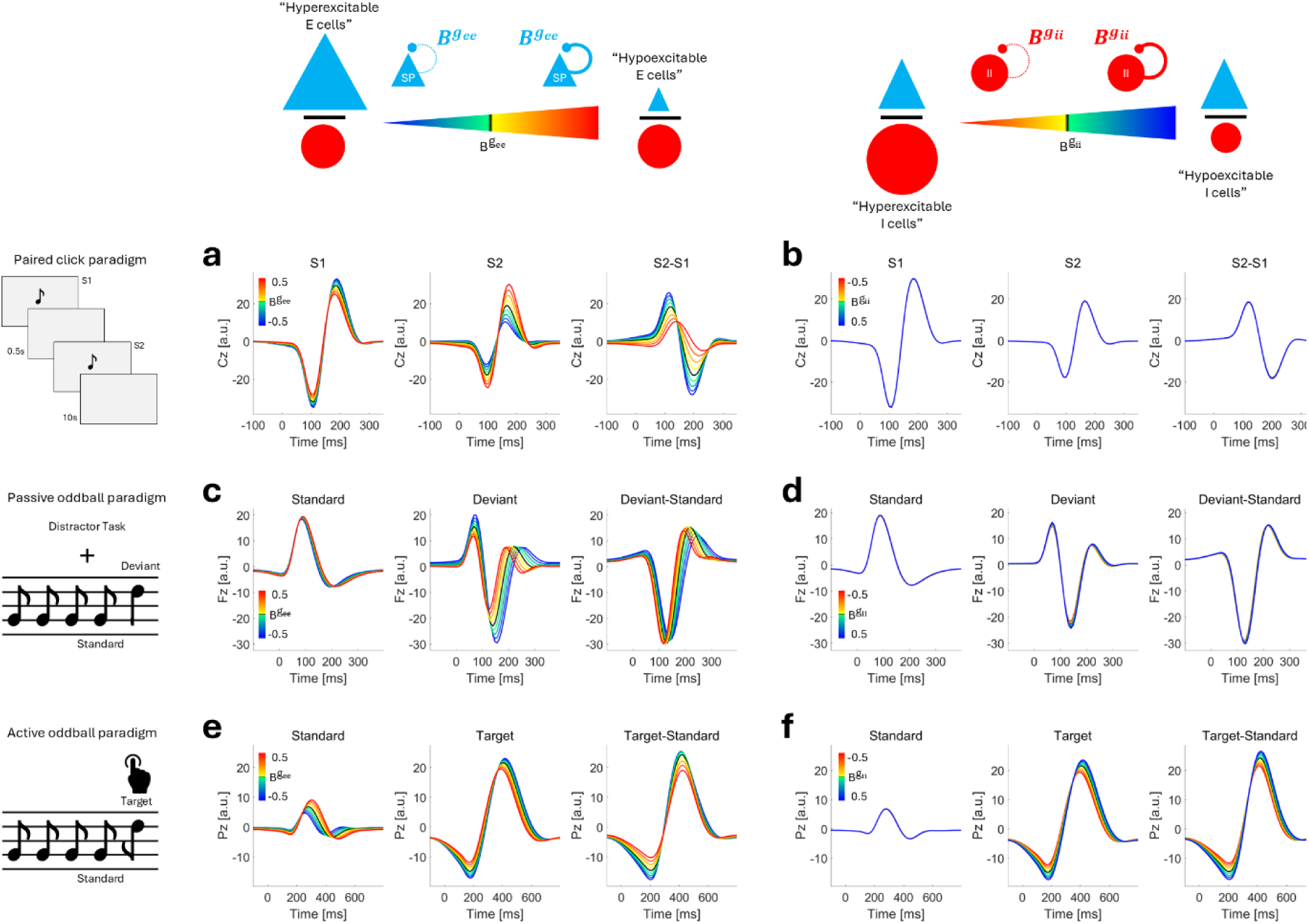
Simulating changes in condition-specific E and I cell excitability. Shown are simulated responses in the paired-click paradigm (**a**, **b**), the passive oddball paradigm (**c**, **d**) or the active oddball paradigm (**e**, **f**) when changing condition-specific E cell excitability *B*^*g_ee_*^ (left) or I cell excitability *B*^*g_ii_*^ (right) over the grid: [-0.5 -0.375 -0.25 -0.125 0 0.125 0.25 0.375 0.5] around the posterior expectation of the parameters (black line). Colder colours indicate increased and warmer colours reduced E/I ratio. Note that increasing the parameter corresponds to a loss of excitability since we parameterised excitability through an inhibitory self-connection.

In the passive oddball paradigm, we found that a loss of E cell excitability (↑*B*^*g_ee_*^) resulted in reduced ERP amplitudes in responses to deviant tones and an overall reduced difference waveform with reduced mismatch negativity and P3a responses (Figure 3c), both observed in schizophrenia (Bramon et al., 2004; Erickson et al., 2016; Hamilton et al., 2019, 2022; Jeon et al., 2003; Umbricht & Krljes, 2005). A similar change in I cell excitability resulted in no comparable change in the ERP (Figure 3d).

For the active oddball paradigm, we observed distinct effects of changing E and I cell excitability. While reduced E/I ratio - either through a loss of E cell excitability (↑*B*^*g_ee_*^) or hyperexcitable I cells (↓*B*^*g_ii_*^) - resulted in reduced P3b in the target-standard difference waveform (Figure 3e), changing E cell excitability affected the ERPs in response to both standard and target tones, whereas changing I cell excitability only affected the response to target tones (Figure 3f). To investigate whether this qualitative distinction would also hold in the passive oddball paradigm, we increased the simulation range for I cell excitability to ±200% (not shown) and indeed observed similar changes (i.e., much stronger effects on the deviant- compared to the standard-ERP). This finding is in line with P3b reductions observed in schizophrenia patients (Bramon et al., 2004; Jeon et al., 2003) and suggests that the model can potentially distinguish E and I cell dysfunction because they relate to different data features.

### Interactions between E and I cell excitability

Next, we investigated whether E and I cell excitability interacted, by systematically varying both parameters together. We found that the effect of changing E cell excitability on the ERP amplitude was largely independent of varying I cell excitability in the paired-click paradigm, as indicated by the straight vertical line in Figure 4a, which suggests that changing E cell excitability has the same effect on the P2 component irrespective of assumed I cell excitability. We observed a similar result for the N2/MMN components in the passive oddball paradigm (Figure 4b). However, we found that this was primarily driven by the effect of changing E cell excitability on the deviant response for all levels of I cell excitability. There was no effect on the standard-ERP. Lastly, we found non-trivial interactions for the P3b component: reducing the E/I ratio through simulating both a loss of E cell excitability (↑*B*^*g_ee_*^) and an increase in I cell excitability (↓*B*^*g_ii_*^) resulted in the most pronounced effects on P3b components of the target-ERP and the difference waveform, whereas the effects of changing E cell excitability on the standard-ERP were largely independent of I cell excitability (Figure 4c). This suggests that P3b amplitudes could reflect changes in both parameters rather than E or I cell function alone (see Supplement for more detailed investigations of parameter interactions for this paradigm).

**Figure 4.**
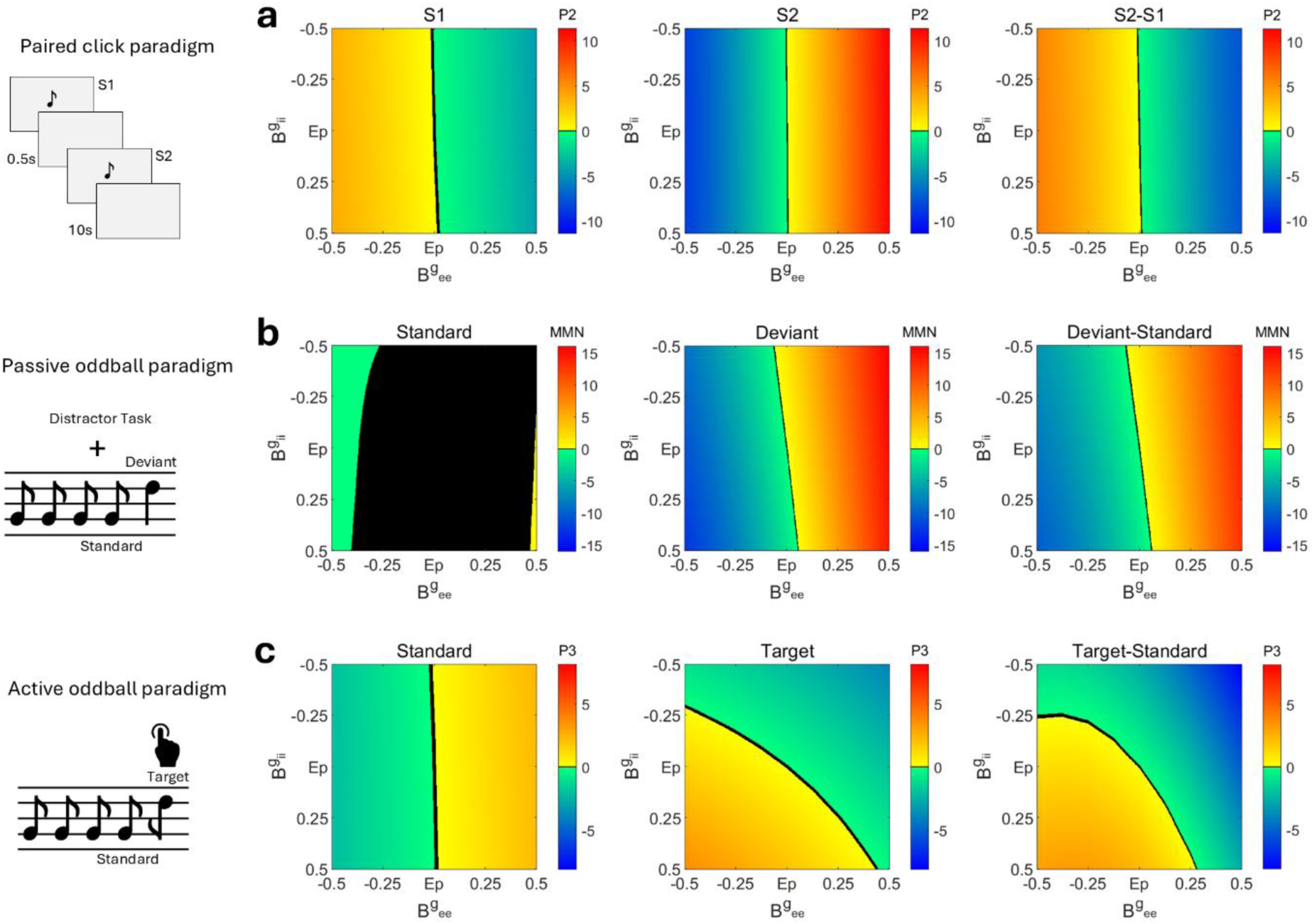
Interactions between E and I cell excitability. We simulated data for changing condition-specific E (*B*^*g_ee_*^) and I (*B*^*g_ii_*^) parameters over the grid: [-0.5 -0.375 -0.25 -0.125 0 0.125 0.25 0.375 0.5]. The remaining values are linearly interpolated for visualization purposes only. To illustrate the effects of changing E/I parameters on the EEG, we selected a characteristic EEG component for each paradigm, specifically the P2 (maximum Pz amplitude between 120-250ms) for the paired-click paradigm (**a**), the mismatch negativity (**MMN**; minimum Fz amplitude between 150-250ms) in the passive oddball paradigm (**b**) and P3 (maximum Pz amplitude between 250-600ms) for the active oddball paradigm (**c**). The black colour indicates EEG component amplitudes that were equivalent to the amplitudes generated with the posterior parameter values. Warmer and colder colours indicate more positive and negative EEG component amplitudes, respectively. All component amplitudes were normalised by subtracting the ERP amplitude generated under the posterior parameter estimates.

## Discussion

This study set out to introduce a canonical microcircuit model to estimate E/I balance from non-invasive M/EEG recordings by parameterising global excitatory and inhibitory cell function. We illustrated how this model can be used to study condition-specific changes in E/I balance by applying it to three different auditory paradigms: paired-click, active oddball and passive oddball. We found that E/I parameters can be recovered efficiently using a variety of empirical paradigms in healthy participants. We then illustrated— with numerical studies—that a loss of pyramidal cell excitability can account for many known empirical findings in the schizophrenia M/EEG literature, including reduced ERP amplitudes across all three paradigms. Lastly, we provide the requisite code, integration schemes and empirical priors for future analyses of data from clinical cohorts.

We found excellent parameter recovery for the parameter capturing pyramidal cell excitability, rendering it a promising and most importantly, biologically interpretable biomarker. This parameter could capture multiple biological processes that change pyramidal cell excitability, for example, NMDA receptor function on pyramidal cells as suggested previously (Adams et al., 2022), or neuromodulatory effects or loss of synapses within the superficial pyramidal cell pool. Interestingly, our simulations suggest that a loss of pyramidal cell excitability may be sufficient to explain the effects observed in patients with schizophrenia across three different auditory paradigms, namely reduced responses in the paired-click, active oddball and passive oddball paradigms (Bramon et al., 2004; Erickson et al., 2016; Hamilton et al., 2019, 2022; Jeon et al., 2003; Umbricht & Krljes, 2005; Turetsky et al., 2008; Shen et al., 2020; Rosburg, 2018).

Secondly, our simulation analysis showed that changing a single model parameter is sufficient to explain changes in the MMN. This provides a very parsimonious explanation without sacrificing biological plausibility compared to previous findings, which reported changes in many different parameters including local excitability as well as extrinsic forward, backward and cross-hemispheric connectivity (Gütlin et al., 2025). Our model is still in line with predictive coding accounts of the MMN (Garrido et al., 2009a; Charlton et al., 2025), which view the MMN as a precision-weighted prediction error signal that is changed in psychosis (Charlton et al., 2022; Hauke et al., 2023). A key feature of our model is that it enables studying E/I balance, predictive coding and their relationship within the same model, unlike other models that have focused on one view or the other, for example by only modelling E and I populations (Deco et al., 2014; Fagerholm et al., 2021). In terms of computational psychiatry, E/I balance is a key mediator of cortical gain control (Abbott et al., 1997), where gain control underwrites belief updating in neuronal circuits based upon predictive coding. Specifically, the synaptic gain of superficial pyramidal cells (generally assumed to encode prediction errors) is read as encoding the precision of prediction errors. A failure of gain control can then be read in terms of functional dysconnections that lead to false inference (Adams et al., 2013; Friston, 2023).

Our model also includes deep pyramidal cells, which are thought to send predictions through extrinsic backward connections (Bastos et al., 2012). The excitability of these cells can be parameterised by adding another global E parameter to their self-inhibition and relaxing the prior on their contribution to the M/EEG signal in the future. This would allow testing whether the excitability of superficial and deep pyramidal cells differently relates to precision-weighted prediction errors and predictions, respectively.

According to the predictive coding view, changing pyramidal cell excitability corresponds to changing the precision-weighting or learning rate used to scale prediction errors elicited by deviant tones. Thus, our results suggest that a failure of precision-weighting can explain reduced ERPs across all three paradigms. Importantly, this scaling likely involves NMDA receptor-related processing (Adams et al., 2022), because cortical gain parameters scale or modulate synaptic connections multiplicatively (Friston et al., 2016; Stephan, Baldeweg, & Friston, 2006) as opposed to additively unlike other cellular processes such as spatial and temporal summation in the cell soma. The contributions of AMPA, GABA and NMDA receptors can be further explored in future studies with the conductance-based version of the canonical microcircuit we provide here.

Beyond schizophrenia, we believe this model can be used to study E/I balance alterations across various other clinical conditions and the effect of pharmacological interventions on E/I. For example, it is well suited for understanding changes in E/I balance in depression (Hu et al., 2023) and the effects of fast-acting antidepressant effects observed following ketamine administration (Hashimoto, 2019; Fagerholm et al., 2021) by modelling suitable M/EEG biomarkers like the MMN. Another interesting avenue for future research will be to apply this model to understand how E/I balance is altered under the influence of psychedelics. Leptourgos et al. (2022) recently proposed that psychedelic-induced hallucinations are likely caused by different circuit changes compared to hallucinations associated with schizophrenia. It has been suggested that 5-HT2A receptor activation on pyramidal cells may lead to disinhibition of these cells (Aghajanian & Marek, 1999; Nichols, 2016).

Our model could be used to test this theory empirically and one may expect to find increased excitability of pyramidal cells following psychedelic administration, which would capture 5-HT2A- mediated disinhibition. Additionally, both predictive coding accounts (Carhart-Harris & Friston, 2019) and E/I balance accounts of psychedelics (Leptourgos et al., 2022; Bedford et al., 2023) have been proposed and can be investigated together. Our model can be used to determine whether the excitability of superficial pyramidal cells associated with prediction errors or deep pyramidal cells sending back predictions are changed under psychedelics as proposed previously (Carhart-Harris & Friston, 2019; Leptourgos et al., 2022). In the future, it will also be important to study E/I imbalance in other disorders including epilepsy (Fritschy, 2001; Rosch et al., 2019; Rosch et al., 2018a; Rosch et al., 2018b), autism (Rubenstein, & Merzenich, 2003) and Alzheimer’s (Maestú et al., 2021; Bi et al., 2020).

There are several limitations of this study that merit attention. First, the recovery for the inhibitory cell excitability parameters was only fair. While this suggests the parameters can in principle be recovered from these paradigms, its recovery will need to be improved, especially for clinical applications. Future work should investigate whether other paradigms like the 40 Hz auditory steady state paradigm (Grent et al., 2023) or resting-state data are better suited for probing inhibitory cell function. Secondly, it will be important to investigate the test-retest reliability of E/I parameters, because our parameter recovery results only provide an upper bound on the reliability of this computational assay of E/I balance (Karvelis, Paulus, & Diaconescu, 2024). Third, since we parameterise E cell excitability through an (inhibitory) self-connection, it is also possible that this parameter reflects a local inhibitory loop perhaps involving parvalbumin-positive (PV+) interneurons. This will need to be clarified in animal studies or by using induced pluripotent stem cell models (iPSCs). iPSCs will also be important to inform priors and constrain parameters (e.g., synaptic time constants) to biologically plausible values in the future.

In conclusion, we introduced a canonical microcircuit for estimating E/I balance. Our simulations suggest that a loss of pyramidal cell excitability may account for various ERP changes observed in schizophrenia. We hope that this tool can serve as a computational assay to assess cell and receptor function based on non-invasive M/EEG recordings (Frässle et al., 2016; Karvelis et al., 2022), enabling the study of E/I balance across various clinical conditions and ultimately permitting allocation of different treatments to individual patients to restore E/I balance and alleviate symptoms.

## Code availability

The posterior parameter estimates (empirical priors for future analyses) and the analysis code for both convolution- and conductance-based models for different integration schemes used in SPM12, TAPAS and the MATLAB default integrator (dde23) will be made publicly available on GitHub upon acceptance of this manuscript (https://github.com/daniel-hauke/dcm_ei).

## Data availability

Data for the paired-click paradigm is freely available through the NIMH data archive (https://nda.nih.gov/edit_collection.html?id=2274). Access to NAPLS2 EEG data can be arranged through a formal collaboration with the NAPLS2 PIs. Contact Dr. Daniel Mathalon to inquire about such collaborations.

## Acknowledgments

We thank the B-SNIP and NAPLS2 consortia for generously providing access to the healthy control data used in this study. The B-SNIP and NAPLS data collection was funded by the NIMH.

