## Supplementary Material for "A Canonical Microcircuit for Estimating Excitation/Inhibition (E/I) Balance"

**Affiliations:**

### Supplementary Methods

#### EEG

B-SNIP EEG data (paired-click paradigm) was recorded through 64 Ag/AGCl sensors (impedance < 5 KΩ; Quik-Cap, Compumedics Neuroscan, El Paso, TX) with a standard 10-10 setup and mastoid electrodes and CP1/2 locations to provide sampling below the canthomeatal line, with nose reference and forehead ground (Hamm et al., 2014). Recordings were amplified (x12,500) and digitized at 1000 Hz using Neuroscan Acquire and Synaps2 recording systems (Compumedics Neuroscan). In NAPLS2, EEG data and vertical-horizontal oculograms were recorded using an Active Two system with mastoid reference electrodes (Biosemi) with 32-channel (4 sites) or 64-channel (4 sites) electrode caps and digitized at 1024 Hz (Hamilton et al., 2019; Hamilton et al., 2022).

#### Preprocessing

Preprocessing of B-SNIP data was conducted using SPM12 (v7771, Wellcome Trust Centre for Neuroimaging, London, UK; https://github.com/spm/spm12/releases/tag/r7771) and EEGLab (version 14.1b, UC San Diego, San Diego, CA, USA; https://sccn.ucsd.edu/eeglab) in Matlab (version 2020a, The MathWorks Inc., Natick, MA, USA; https://mathworks.com). Data from NAPLS2 were preprocessed using EEGLab (UC San Diego, San Diego, CA, USA; https://sccn.ucsd.edu/eeglab) in Matlab (The MathWorks Inc., Natick, MA, USA; https://mathworks.com), as described in Hamilton et al. (2019, 2022).

##### Paired-click paradigm

Preprocessing followed the steps described in Adams et al. (2022). Data was imported into EEGlab, average-referenced, high-pass filtered (1 Hz), notch-filtered (59.5-60.5Hz), low-pass (70 Hz) and epoched from -100 ms to 400 ms around tone onsets. Subsequently, we ran an automated artefact detection pipeline, which involved the following steps. Trials were classified as bad trials, if the SD of their normalised amplitude fell outside of [-5 5], if they included linear trends identified with pop_rejtrend (max slope=100, min R2=0.8), based on frequency thresholds with pop_rejspec.m (0-2 Hz, [-50 dB 50 dB]; 20-40Hz, [-100 dB 25 dB]), based on joint probability thresholding with pop_jointprob.m or kurtosis with pop_rejkurt.m (local SD threshold: 5, global SD threshold: 4 for both). If this procedure rejected 50% of the trials then bad channel rejection was run first. This involved rejecting bad channels with pop_rejchan.m based on extreme values in their spectrum ([-4 6] SD), kurtosis ([-7 12]), joint probability ([-8 7]). If there were still >50% bad trials or >20% bad channels the dataset was discarded. Next, we performed independent component analysis (ICA) using the Second Order Blind Identification (SOBI) algorithm and components that were automatically classified as artefacts with p>0.5 – determined using the Multiple Artefact Rejection Algorithm (MARA; Winkler et al., 2011) – were then rejected. MARA is a supervised machine learning algorithm trained based on expert ratings of 1290 artefact components (e.g., eye, muscle, loose electrode) to identify artefact components based on spectral, spatial and temporal features. Following MARA, we repeated bad channel and epoch rejection as some artefacts only became apparent after ICA. Bad channels were identified based on extreme values in their spectrum ([-6 5] SD), kurtosis ([-6 9]), joint probability ([-7 7]). Bad epochs were rejected based on joint probability and kurtosis (local SD threshold: 4, global SD threshold: 3 for both). Bad channels were interpolated. Subsequently, data was imported into SPM, baseline corrected (-100 ms to -50 ms), downsampled to 250 Hz and averaged using SPM's robust averaging. For DCM analysis, we additionally performed a post hoc low-pass filtering step (30 Hz) to remove high-frequency noise.

##### Passive oddball paradigm

Data were re-referenced to a mastoid average, high-pass filtered (1Hz) and segmented into 1s epochs. Eyeblink correction was performed using a regression method (Gratton et al., 1983) and epochs were baseline corrected (-50 to 0 ms). Spherical spline interpolation was performed (Delorme

& Makeig, 2004) and epochs were removed as bad trials if their amplitude exceeded ±100 µV. Averages were computed using sorted averaging (Rahne et al., 2008). For more details, please see Hamilton et al. (2022). Preprocessed data were imported into SPM and downsampled to 250 Hz. Grand mean waveforms were computed using the 32 channels common to everyone, while the parameter identifiability and recovery analyses retained the original number of channels from each participant's data.

##### Active oddball paradigm

Data were re-referenced to a mastoid average, high-pass filtered (0.1Hz) and segmented into 3s epochs (-1s to 2s around tone onsets). Artefact rejection was conducted using fully automated statistical thresholding (Nolan, Whelan, & Reilly, 2010) and canonical correlation analysis (De Clercq et al., 2006; Riès et al., 2013). Lastly, epochs were baseline corrected (-100 to 0 ms). For more details, please see Hamilton et al. (2019). Preprocessed data were imported into SPM and downsampled to 250Hz. For DCM analysis, we additionally performed a post hoc low-pass filtering step (15 Hz) to remove high-frequency noise. Grand mean waveforms were computed using the 32 channels common to everyone, while the parameter identifiability and recovery analyses retained the original number of channels from each participant's data.

#### Source reconstruction

Although source reconstructions are not recommended for 32-channel data, please note that we do not optimise the source locations in our DCM model, but rather keep them fixed based on previous literature, which makes the inverse problem tractable. Instead, the focus of our approach is to understand the neuronal dynamics that give rise to the EEG signals. The source localisation analysis in the main manuscript should just be considered a sanity check to confirm that we can indeed explain sufficient variance with the selected network.

#### Integration

We integrated the convolution-based model with a Euler integration scheme based on Schöbi et al. (2021). This integrator reduces integration errors that can be introduced through the absorption of delay operators into the Jacobian and a linear approximation to the delayed neuronal states, which takes place while integrating delay differential equations with the default SPM12 integrator for the original Jansen & Rit model (spm_int_L). Under some circumstances (e.g., long delays) these integration errors can reach amplitudes similar to the magnitude of the ERP signals (Schöbi et al., 2021). Note, however, that SPM12 also includes more accurate integration schemes that approximate the delays using an *n*-th order polynomial (spm_dcm_delay), which is the default for the CMC model, or options to re-evaluate the Jacobian in each iteration (spm_int_J). However, these more accurate integrators also come with a computational cost paid in runtime. In practice, we found that the Euler integrator provided the best trade-off between accuracy and computational cost for our purposes. The integrator code is openly available as part of the Translational Algorithms for Psychiatry-Advancing Science's code toolbox (TAPAS version R09.2020; Frässle et al., 2021; https://www.translationalneuromodelling.org/tapas).

For the conductance-based model, we provide code that works with different integration schemes, recognising that the choice of the integration scheme depends on the computational resources available and the accuracy required for the specific application in mind. Specifically, we provide code that works with all the default SPM integrators, as well as an additional implementation using MATLAB's default dde23 integrator, which employs a more accurate, but much slower, Runge-Kutta based method (Shampine, 2001). The dde23 integrator allows users to specify integration error bounds and is arbitrarily accurate, provided that users are willing to run their model for a long time. It can be used for benchmarking other integration schemes, as done in previous work (Lemaréchal et al., 2018; Schöbi, 2020; Schöbi et al., 2021; Do Tri, 2022).

### Supplementary Results

| Parameter | Description | Posterior Mean | | | | | | Posterior Precision | | | | | |
|---|---|---|---|---|---|---|---|---|---|---|---|---|---|
| **Neuronal Source Model f(x)** | | | | | | | | | | | | | |
| **A{1}** | Average extrinsic connectivity (forward, SP to SS) | | -0.0461 | | | | | | 0.0624 | | | | |
| | | 0.0058 | | | | | | 0.0625 | | | | | |
| | | 0.8094 | | | 0.0072 | | | 0.0546 | | | 0.0625 | | |
| | | | 1.7572 | 0.0102 | | | | | 0.0447 | 0.0624 | | | |
| | | | | 0.6213 | | | 0.0391 | | | 0.0544 | | | 0.0625 |
| | | | | | 1.0074 | -0.0126 | | | | | 0.0521 | 0.0625 | |
| **A{2}** | Average extrinsic connectivity (forward, SP to DP) | | 0.0005 | | | | | | 0.0625 | | | | |
| | | 0 | | | | | | 0.0625 | | | | | |
| | | 0.0084 | | | 0.0001 | | | 0.0625 | | | 0.0625 | | |
| | | | -0.0081 | 0 | | | | | 0.0625 | 0.0625 | | | |
| | | | | -0.0008 | | | -0.0003 | | | 0.0625 | | | 0.0625 |
| | | | | | -0.0225 | 0.0002 | | | | | 0.0625 | 0.0625 | |
| **A{3}** | Average extrinsic connectivity (backward, DP to SP) | | | 0.0665 | | | | | | 0.0623 | | | |
| | | | | | -0.0075 | | | | | | 0.0625 | | |
| | | | | | | 0.0055 | | | | | | 0.0624 | |
| | | | | | | | -0.0440 | | | | | | 0.0625 |
| **A{4}** | Average extrinsic connectivity (backward, DP to II) | | | -0.0094 | | | | | | 0.0625 | | | |
| | | | | | 0.0005 | | | | | | 0.0625 | | |
| | | | | | | -0.0012 | | | | | | 0.0625 | |
| | | | | | | | 0.0063 | | | | | | 0.0625 |
| **B** | Condition-specific extrinsic connectivity | | | 0.1123 | | | | | | 0.1245 | | | |
| | | | | | 0.0486 | | | | | | 0.1250 | | |
| | | 1.5090 | | | | 0.0090 | | 0.0929 | | | | 0.1249 | |
| | | | -0.5429 | | | | 0.0760 | | 0.0063 | | | | 0.1249 |
| | | | | 1.2334 | | | | | | 0.0928 | | | |
| | | | | | -1.9057 | | | | | | 0.0833 | | |
| **C** | Input Strength | -0.4096 | -0.0595 | | | | | 0.0197 | 0.0153 | | | | |
| **G** | Average intrinsic connectivity $[g_{ee}\ g_{ii}]$ | | | 0.0944 | 0.0253 | | | | | 0.0243 | 0.0312 | | |
| **$B^G$** | Condition-specific intrinsic connectivity $[B^{gee}\ B^{gii}]$ | | | -0.8600 | -0.0954 | | | | | 0.0124 | 0.1248 | | |
| **T** | Synaptic time constant [SS SP II DP] ms | | 19.0974 | 11.7137 | 1.3658 | 16.1249 | | | 0.0039 | 0.0081 | 0.0296 | 0.0307 | |
| **D** | Neuronal delays (all sources) [intrinsic extrinsic] ms | | | 1.0890 | 6.5191 | | | | | 0.0154 | 0.0134 | | |
| **S** | Slope of output firing rate function | -1.5639 | | | | | | 0.0022 | | | | | |
| **R** | Stimulus [onset dispersion] ms | | | 68.4704 | 20.3564 | | | | | 0.0001 | 0.0006 | | |
| **Forward Model g(x)** | | | | | | | | | | | | | |
| **Lpos** | Source location [MNI coordinates] | -53 | -20 | 13 | | | | 0 | | | | | |
| | | 49 | -10 | 10 | | | | | | | | | |
| | | -36 | 23 | 3 | | | | | | | | | |
| | | 36 | 23 | 3 | | | | | | | | | |
| | | -1 | -20 | 67 | | | | | | | | | |
| | | 1 | -20 | 67 | | | | | | | | | |
| **L** | Dipole orientation | 2.1949 | -0.7451 | -0.0195 | -6.0513 | 2.8187 | -1.6533 | 0.0938 | 0.0220 | 0.1828 | 2.3506 | 1.6952 | 2.2203 |
| | | -3.0705 | 0.0387 | 3.0591 | 0.8621 | 21.0115 | 24.3045 | 0.1506 | 0.0247 | 1.2119 | 1.6964 | 24.4919 | 35.5514 |
| | | -4.4568 | -3.5255 | 5.2690 | 18.2303 | 24.3745 | 16.3025 | 0.3268 | 0.1556 | 4.0038 | 17.3483 | 32.0612 | 17.4809 |
| **J** | Neuronal populations contributing to EEG: [SS SP II DP] | 0 | 1 | 0 | 0 | | | 0 | | | | | |

**Table S1. Parameter Posteriors for the Paired-Click Paradigm. DP** deep pyramidal cells. **II** inhibitory interneurons. **SP** superficial pyramidal cells. **SS** spiny stellate cells.

| Parameter | Description | Posterior Mean | | | | | | Posterior Precision | | | | | |
|---|---|---|---|---|---|---|---|---|---|---|---|---|---|
| **Neuronal Source Model f(x)** | | | | | | | | | | | | | |
| **A{1}** | Average extrinsic connectivity (forward, SP to SS) | | 0.1660 | | | | | | 0.0620 | | | | |
| | | 0.0049 | | | | | | 0.0622 | | | | | |
| | | -0.0028 | | | 0.0302 | | | 0.0537 | | | 0.0623 | | |
| | | | 0.1283 | 0.0299 | | | | | 0.0534 | 0.0623 | | | |
| | | | | 0.2888 | | | -0.0384 | | | 0.0542 | | | 0.0624 |
| | | | | | 0.2838 | -0.0185 | | | | | 0.0544 | 0.0623 | |
| **A{2}** | Average extrinsic connectivity (forward, SP to DP) | | -0.0006 | | | | | | 0.0625 | | | | |
| | | 0 | | | | | | 0.0625 | | | | | |
| | | -0.0006 | | | -0.0001 | | | 0.0625 | | | 0.0625 | | |
| | | | -0.0004 | 0 | | | | | 0.0625 | 0.0625 | | | |
| | | | | -0.0009 | | | 0.0001 | | | 0.0625 | | | 0.0625 |
| | | | | | 0.0011 | 0 | | | | | 0.0625 | 0.0625 | |
| **A{3}** | Average extrinsic connectivity (backward, DP to SP) | | | -0.0028 | | | | | | 0.0625 | | | |
| | | | | | -0.0091 | | | | | | 0.0625 | | |
| | | | | | | -0.0031 | | | | | | 0.0625 | |
| | | | | | | | -0.0057 | | | | | | 0.0625 |
| **A{4}** | Average extrinsic connectivity (backward, DP to II) | | | -0.0003 | | | | | | 0.0625 | | | |
| | | | | | 0.0016 | | | | | | 0.0625 | | |
| | | | | | | 0.0007 | | | | | | 0.0625 | |
| | | | | | | | 0.0014 | | | | | | 0.0625 |
| **B** | Condition-specific extrinsic connectivity | | | -0.0098 | | | | | | 0.1249 | | | |
| | | | | | -0.0198 | | | | | | 0.1249 | | |
| | | 1.8700 | | | | -0.0023 | | 0.0588 | | | | 0.1250 | |
| | | | 1.6228 | | | | -0.0061 | | 0.0365 | | | | 0.1250 |
| | | | | 0.6810 | | | | | | 0.0925 | | | |
| | | | | | 0.7033 | | | | | | 0.0926 | | |
| **C** | Input Strength | -0.0313 | -0.0030 | | | | | 0.0281 | 0.0286 | | | | |
| **G** | Average intrinsic connectivity $[g_{ee}\ g_{ii}]$ | -0.9173 | 0.3557 | | | | | 0.0214 | 0.0306 | | | | |
| **$B^G$** | Condition-specific intrinsic connectivity $[B^{gee}\ B^{gii}]$ | 0.9197 | 1.4871 | | | | | 0.0120 | 0.1093 | | | | |
| **T** | Synaptic time constant [SS SP II DP] ms | 8.8903 | 54.1230 | 2.3014 | 1.8839 | | | 0.0173 | 0.0008 | 0.0224 | 0.0312 | | |
| **D** | Neuronal delays (all sources) [intrinsic extrinsic] ms | -0.0004 | -0.0385 | | | | | 0.0155 | 0.0137 | | | | |
| **S** | Slope of output firing rate function | -1.0497 | | | | | | 0.0043 | | | | | |
| **R** | Stimulus [onset dispersion] ms | 126.6261 | 15.3041 | | | | | 0.0001 | 0.0008 | | | | |
| **Forward Model g(x)** | | | | | | | | | | | | | |
| **Lpos** | Source location [MNI coordinates] | -42 | -22 | 7 | | | | 0 | | | | | |
| | | 46 | -14 | 8 | | | | | | | | | |
| | | -61 | -32 | 8 | | | | | | | | | |
| | | 59 | -25 | 8 | | | | | | | | | |
| | | -46 | 20 | 8 | | | | | | | | | |
| | | 46 | 20 | 8 | | | | | | | | | |
| **L** | Dipole orientation | 0.5687 | -0.2568 | 3.0594 | -4.7348 | 11.8960 | -19.0720 | 0.0189 | 0.0127 | 0.9936 | 2.1980 | 13.3038 | 36.2121 |
| | | 1.6366 | 1.2313 | -3.4501 | -4.4330 | 8.5635 | 1.0112 | 0.0679 | 0.0432 | 1.2416 | 1.9413 | 9.4963 | 4.0926 |
| | | 1.2269 | 0.0851 | -5.5316 | -2.5471 | -21.0412 | -9.2916 | 0.0530 | 0.0185 | 3.1236 | 0.7947 | 35.2698 | 13.2876 |
| **J** | Neuronal populations contributing to EEG: [SS SP II DP] | 0 | 1 | 0 | 0 | | | 0 | | | | | |

**Table S2. Parameter Posteriors for the Passive Oddball Paradigm. DP** deep pyramidal cells. **II** inhibitory interneurons. **SP** superficial pyramidal cells. **SS** spiny stellate cells.

| Parameter | Description | Posterior Mean | | | | | | Posterior Precision | | | | | |
|---|---|---|---|---|---|---|---|---|---|---|---|---|---|
| **Neuronal Source Model f(x)** | | | | | | | | | | | | | |
| **A{1}** | Average extrinsic connectivity (forward, SP to SS) | | 0.1315 | | | | | | 0.0618 | | | | |
| | | 0.3525 | | | | | | 0.0613 | | | | | |
| | | 0.6031 | | | 1.5925 | | | 0.0404 | | | 0.0413 | | |
| | | | 0.2476 | 2.2975 | | | | | 0.0400 | 0.0357 | | | |
| | | | | -0.0852 | | | 0.0361 | | | 0.0477 | | | 0.0609 |
| | | | | | -0.2897 | 0.0202 | | | | | 0.0515 | 0.0610 | |
| **A{2}** | Average extrinsic connectivity (forward, SP to DP) | | -0.0006 | | | | | | 0.0625 | | | | |
| | | -0.0003 | | | | | | 0.0625 | | | | | |
| | | 0.0006 | | | -0.0010 | | | 0.0625 | | | 0.0625 | | |
| | | | -0.0028 | -0.0004 | | | | | 0.0625 | 0.0625 | | | |
| | | | | -0.0050 | | | -0.0007 | | | 0.0625 | | | 0.0625 |
| | | | | | 0.0001 | -0.0002 | | | | | 0.0625 | 0.0625 | |
| **A{3}** | Average extrinsic connectivity (backward, DP to SP) | | | -0.0508 | | | | | | 0.0625 | | | |
| | | | | | -0.0199 | | | | | | 0.0625 | | |
| | | | | | | -0.0355 | | | | | | 0.0625 | |
| | | | | | | | -0.0141 | | | | | | 0.0625 |
| **A{4}** | Average extrinsic connectivity (backward, DP to II) | | | 0.0067 | | | | | | 0.0625 | | | |
| | | | | | 0.0029 | | | | | | 0.0625 | | |
| | | | | | | 0.0046 | | | | | | 0.0625 | |
| | | | | | | | 0.0025 | | | | | | 0.0625 |
| **B** | Condition-specific extrinsic connectivity | | | -0.0788 | | | | | | 0.1249 | | | |
| | | | | | -0.0278 | | | | | | 0.1249 | | |
| | | -0.0529 | | | | -0.0549 | | 0.0033 | | | | 0.1250 | |
| | | | -0.0827 | | | | -0.0176 | | 0.0052 | | | | 0.1250 |
| | | | | -0.0975 | | | | | | 0.0102 | | | |
| | | | | | -0.2822 | | | | | | 0.0179 | | |
| **C** | Input Strength | -0.2726 | -0.3930 | | | | | 0.0267 | 0.0264 | | | | |
| **G** | Average intrinsic connectivity [$g_{ee}$ $g_{ii}$] | | | -1.6852 | 0.5987 | | | | | 0.0219 | 0.0294 | | |
| **$B^G$** | Condition-specific intrinsic connectivity [$B^{gee}$ $B^{gii}$] | | | -0.09779 | 2.3360 | | | | | 0.0251 | 0.0957 | | |
| **T** | Synaptic time constant [SS SP II DP] ms | | 21.0609 | 65.3824 | 3.0517 | 1.7659 | | | 0.0153 | 0.0021 | 0.0253 | 0.0312 | |
| **D** | Neuronal delays (all sources) [intrinsic extrinsic] ms | | | -0.0003 | -0.0681 | | | | | 0.0156 | 0.0149 | | |
| **S** | Slope of output firing rate function | -1.3414 | | | | | | 0.0050 | | | | | |
| **R** | Stimulus [onset dispersion] ms | | | 198.1853 | 15.5295 | | | | | 0.0004 | 0.0010 | | |
| **Forward Model g(x)** | | | | | | | | | | | | | |
| **Lpos** | Source location [MNI coordinates] | | | -61 | -32 | 8 | | 0 | | | | | |
| | | | | 59 | -25 | 8 | | | | | | | |
| | | | | -56 | 7 | 29 | | | | | | | |
| | | | | 50 | 8 | 30 | | | | | | | |
| | | | | -33 | -42 | 64 | | | | | | | |
| | | | | 33 | -42 | 64 | | | | | | | |
| **L** | Dipole orientation | 0.2559 | -0.5684 | 1.4549 | -0.0322 | -12.6200 | 17.5433 | 0.0036 | 0.0105 | 0.1066 | 0.0335 | 11.3082 | 23.9110 |
| | | 0.3624 | 0.2780 | -2.0442 | -3.1267 | -17.5307 | -4.2776 | 0.0051 | 0.0043 | 0.2141 | 0.5021 | 21.8944 | 3.9224 |
| | | 0.0702 | 0.0365 | 1.1899 | -0.6768 | -5.3776 | 6.4451 | 0.0039 | 0.0047 | 0.0892 | 0.0655 | 2.5536 | 4.4849 |
| **J** | Neuronal populations contributing to EEG: [SS SP II DP] | | | 0 | 1 | 0 | 0 | 0 | | | | | |

**Table S3. Parameter Posteriors for the Active Oddball Paradigm. DP** deep pyramidal cells. **II** inhibitory interneurons. **SP** superficial pyramidal cells. **SS** spiny stellate cells.

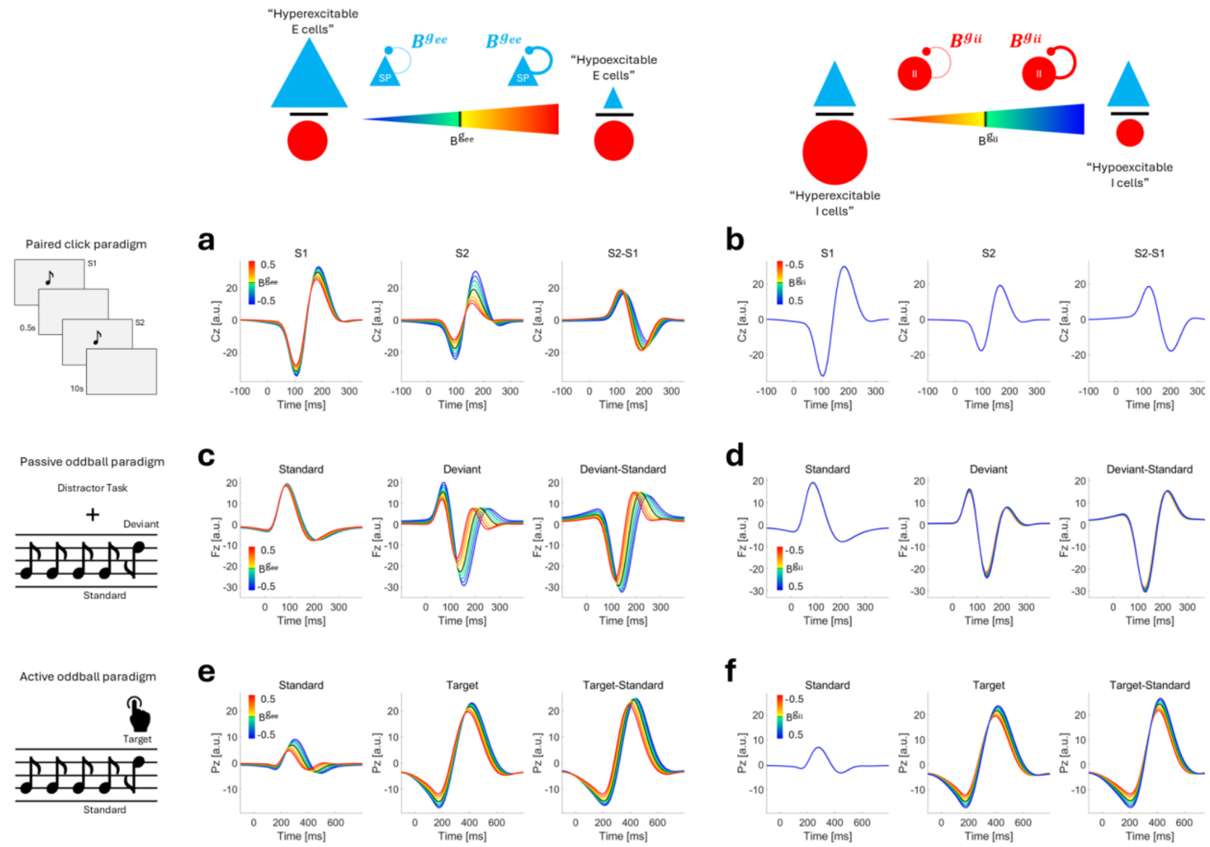

Figure S1. **Simulating changes in condition-independent E and I cell excitability.** Shown are simulated responses in the paired-click paradigm (**a**, **b**), the passive oddball paradigm (**c**, **d**) or the active oddball paradigm (**e**, **f**) when changing condition-independent baseline E cell excitability $g_{ee}$ (left) or I cell excitability $g_{ii}$ (right) over the grid: [-0.5 -0.375 -0.25 -0.125 0 0.125 0.25 0.375 0.5] around the posterior expectation of the parameters (black line). Colder colours indicate increased and warmer colours reduced E/I ratio. Note that increasing the parameter corresponds to a loss of excitability since we parameterised excitability through an inhibitory self-connection.

#### Interactions with other parameters

Since we observed an interaction between E/I parameters in the active oddball paradigm, we investigated their relationships with other parameters that could influence these E/I measures. Specifically, we examined: (1) interactions between condition-specific and condition-independent E/I parameters, and (2) interactions between condition-specific E/I parameters and the time constants of their respective neural populations (Figure S2).

These simulations suggest that the strongest P3b amplitude reductions occur when the E/I ratio of either condition-specific (Figure S2a) or condition-independent E/I parameters (Figure S2d) is reduced, i.e. when there is a loss of E cell excitability combined with hyperexcitable I cells. When comparing the effects of varying both the condition-specific and condition-independent E parameters, the simulations suggest that increasing the condition-specific parameter always reduces the P3b waveform, but under low—but not high—levels of condition-independent connectivity, a condition-specific E cell hyperexcitability (↓$B^{g_{ee}}$) can result in increased P3b responses (Figure S2b). Condition-specific interneuron hyperexcitability leads to reduced P3b responses across different levels of condition-independent I cell excitability suggesting an additive interaction (Figure S2e).

Interestingly, the effects of condition-specific E/I parameters appear to be largely independent of the time-constants of their respective neuronal populations (superior pyramidal cells and interneurons), e.g., hyperexcitable interneurons reduced the P3b amplitude similarly for all parameter values of the inhibitory time constants (Figure S2f). However, for E cells, the time-constant appears to lie in a range that maximises the effect that changing condition-specific pyramidal cell excitability has on the P3b amplitude. Together, these simulations highlight that there can be non-trivial interactions between different model parameters in some of the modelled paradigms that need to be considered in future analyses.

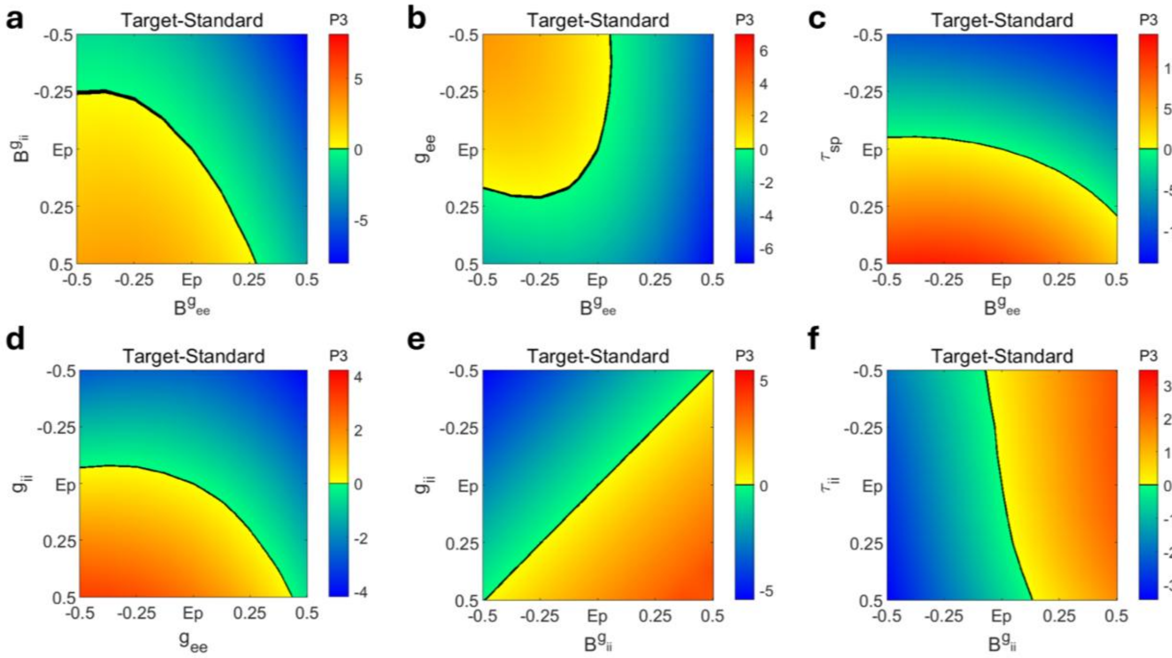

Figure S2**. Assessing interactions with other parameters.** To illustrate interactions between different parameters, we simulated data changing condition-specific ($B^{g_{ee}}$, $B^{g_{ii}}$) (**a**) and condition-independent ($g_{ee}$, $g_{ii}$) (**d**) E/I parameters as well as the time-constants of their respective neuronal populations ($\tau_{sp}$, $\tau_{ii}$) (**b, c, e, f**) over the grid: [-0.5 -0.375 -0.25 -0.125 0 0.125 0.25 0.375 0.5]. The remaining values are linearly interpolated for visualization purposes only. For this analysis, we focus on the active oddball paradigm and plot the change in P3b amplitude of the target-standard difference waveform (maximum Pz amplitude between 250-600ms) normalised by subtracting the P300 amplitude generated under the posterior parameter estimates. The black colour indicates EEG component amplitudes that were close to the amplitudes generated under the posterior. Warmer and colder colours indicate more positive and negative P3b amplitudes, respectively.
